# Aging reduces the number and function of L-type calcium channels in the membrane of cardiac pacemaker cells

**DOI:** 10.1101/2022.06.22.497267

**Authors:** Oscar Vivas, Matthias Baudot, Roxanne Madden, Sabrina Choi, Victor A. Flores, L. Fernando Santana, Claudia M. Moreno

**Author notes:** Corresponding author: Claudia Moreno. Co-first authors (Oscar Vivas, Matthias Baudot).

## Abstract

Every heartbeat is initiated by a spontaneous electrical signal generated inside the cardiac pacemaker. The generation of this electrical signal depends on the coordinated opening and closing of different ion channels, where voltage-gated L-type calcium channels play a central role. Despite the reliability of the pacemaker, all mammals experience a linear slowdown of the pacemaker rate with age. In humans, this slowing can become pathological and constitutes the main cause for the requirement of the implantation of artificial pacemakers. However, the mechanisms behind the age-associated slowdown of the pacemaker are not well understood. Here, we show that age alters L-type calcium channels in pacemaker cells from mice. The age-associated alterations include: i) a reduction in the density of the channels at the plasma membrane, ii) a reduction in the clustering of the channels, and iii) a decrease in channel open probability. Altogether, these age-associated alterations result in a global reduction of the L-type calcium current density and in a slowdown of the pacemaker diastolic depolarization. Remarkably, increasing the open probability of L-type calcium channels pharmacologically was enough to restore pacemaker rate in old cells to the same levels observed in the young. Overall, our findings provide evidence that proper organization and function of L-type calcium channels is impaired by aging and that this dysfunction contributes to the slowdown of pacemaker cells in old animals.

## Introduction

Mammals, including humans and mice, experience a linear decline in heart rate with age^1–5^. How fast the heart beats depends on the activity of the cardiac pacemaker, a specialized region of the heart that fires spontaneous and rhythmic action potentials. The electrical signal originating inside the pacemaker propagates in the form of a calcium wave through the electrical conduction system of the heart, triggering the depolarization and contraction of the atrial and ventricular chambers. This cycle is repeated without interruption from embryonic development until death. The intrinsic pacemaker rate declines linearly from birth at a rate of ∼0.8 bpm/year in humans and ∼4 bpm/month in mice^1^. This natural slowdown of the pacemaker can become pathological. In fact, the age-associated slowdown of the pacemaker is the main cause for the more than half a million artificial pacemaker devices implanted annually in the world^6,7^. However, the molecular mechanisms behind the slowdown of the pacemaker are not completely understood.

The automaticity of the pacemaker relies on a fine balance of ionic currents that drive and allow the regeneration of the pacemaker action potential, where voltage-gated L-type calcium channels play a key role. The pacemaking mechanism can be seen as an oscillatory cycle of the membrane potential. Although oscillatory in nature, the cycle could be described starting in the diastolic depolarization (DD) phase, which can be divided in Early and Late DD. The Early DD involves the activation of hyperpolarization-activated, cyclic nucleotide–sensitive (HCN) channels^8,9^ and T-type Ca_V_3.1 calcium channels^10^ at voltages below -60 mV. The subsequent Late DD starts at around -55 mV with the opening of L-type Ca_V_1.3 channels^10^. This inward current depolarizes the membrane further enough to activate Ca_V_1.2 L-type calcium channels at around -40 mV, pushing the membrane potential to the required threshold to trigger the action potential. The pacemaker action potential is a calcium action potential completely carried by the Ca_V_1.2 and Ca_V_1.3 L-type calcium channels. The cycle is completed by the inactivation of calcium channels and the activation of several potassium channels, including I_to_, I_Kur_, I_Kr_, and I_Ks_, resulting in the repolarization phase of the action potential^11^. Repolarization leads to the activation of the diastolic depolarization, which starts the cycle again.

L-type calcium channels play an essential role not only in the generation of each action potential but also in the control of firing rate. Knockout mice lacking the Ca_V_1.3 channel exhibit bradycardia and sinus pauses^12,13^. In addition, humans carrying point mutations that reduce the function of Ca_V_1.3 channels also exhibit bradycardia^14,15^, resembling the pacemaker dysfunction observed in the elderly. The remodeling of ionic currents, including the down-regulation or up-regulation of the expression of specific ion channels, has been proposed to be one of the primary mechanisms underlying the age-associated dysfunction of the cardiac pacemaker^16^. The best-understood change is the alteration of HCN4 channels. HCN4 current density is reduced, and the voltage-dependent activation is shifted in aged animals^1,17,18^. The age-associated alteration of HCN4 is not controversial. However, how aging alters L-type calcium channels is still under debate.

Support and opposition for age-associated changes in L-type calcium channels have been documented. Ca_V_1.2 mRNA increases slightly, while Ca_V_1.3 mRNA does not change in aged rats^16^. At the protein level, the expression of Ca_V_1.2 channels in the pacemaker has been reported to be reduced by about 40% in aged rats^19^, and almost 80% in aged guinea pigs^20^. Furthermore, the current flowing through L-type calcium channels, which measures the function of the proteins, is reduced in aged mice^1^. In another turn, an in-silico model concluded that the remodeling of I_CaL_, either via a gain or loss of function, is predicted to be the most significant factor underlying the slowing down of cardiac pacemaking rates in old animals^21^. Discrepancies between these studies may relate to different approaches, animal models, or ages. But it merits further revision. Importantly, if the expression or function of these channels is altered with aging, what is the mechanism underlying such alterations?

To contribute to answering this question and reaching a consensus, we combined electrophysiology, microscopy, and pharmacology to study L-type calcium channel function, expression, and organization in pacemaker cells isolated from young and old mice. Our results provide evidence that aging leads to a reduction in L-type calcium channel expression at the plasma membrane, contributing to the pacemaker’s age-associated slowdown.

## Results

### Sinoatrial node and pacemaker cell identification

In this work, we studied the sinoatrial pacemaker, a region delimited at the top by the superior vena cava, at the right by the sulcus terminalis in the right atrium, and at the bottom by the coronary sulcus and the inferior vena cava (Figure 1A). The sinoatrial node artery was also used as an anatomical landmark to identify the pacemaker region.

**Figure 1.**
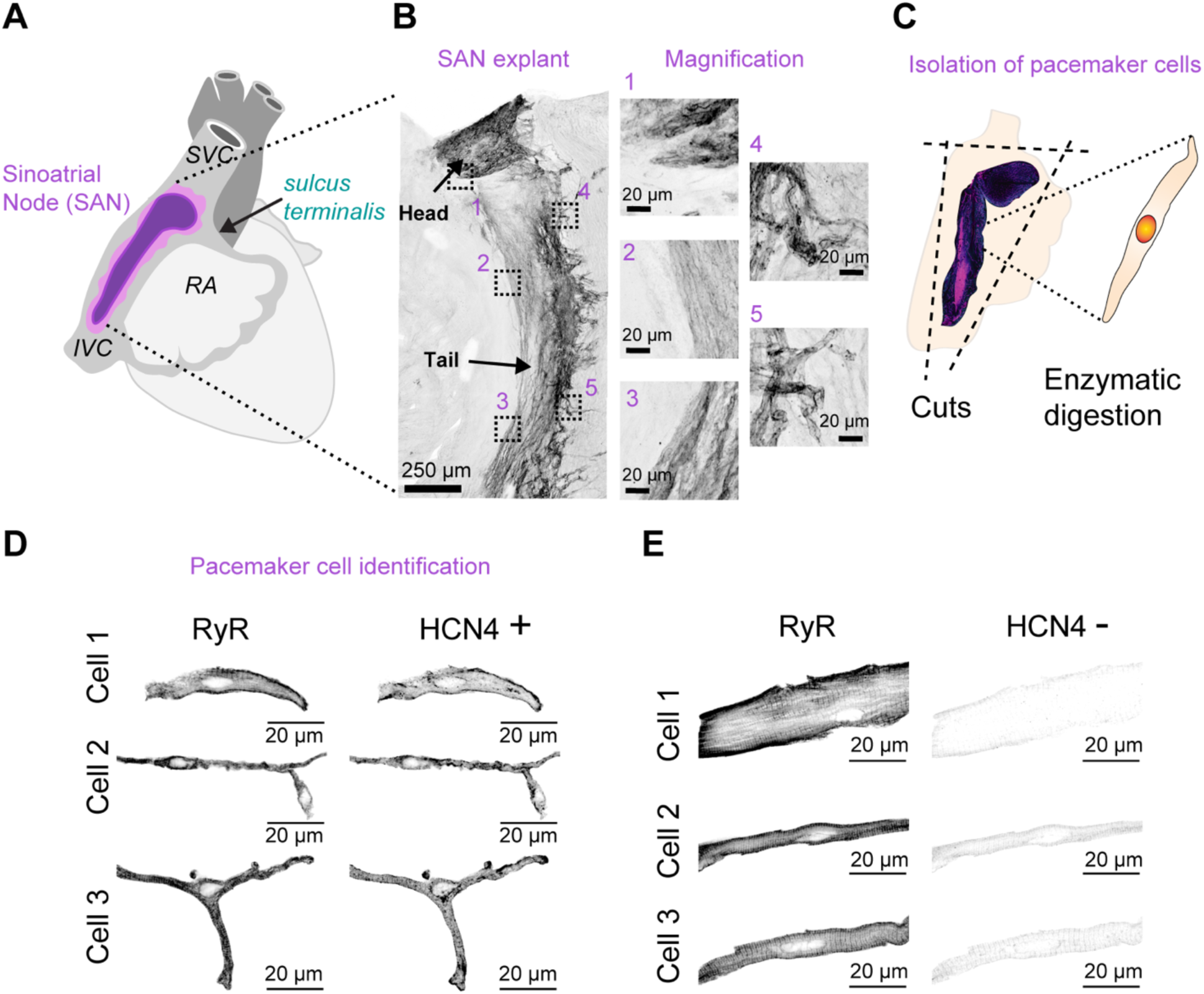
Identification of the sinoatrial node and pacemaker cells. **A.** Schematic representation of the position of the sinoatrial node (SAN) pacemaker in the heart, located between the right atrium (RA) and the superior and inferior vena cava (SVC, IVC). **B.** Volumetric reconstruction of a cleared SAN explant immuno-stained against HCN4 with identified regions for the head and tail. (Right) Magnification of five different regions showing cellular organization inside the tissue. **C.** Schematic representation of the procedure to dissect and isolate single pacemaker cells. **D, E.** Pictures of single cells stained against HCN4 and RyR2 after isolation. Panel D shows three representative pacemaker cells positive for HCN4. Panel E shows three representative cells negative for HCN4.

Explants of this region were immunostained against HCN4, a marker of the pacemaker region. As shown in Figure 1B, the sinoatrial pacemaker is formed by cells organized into defined head and tail structures. For live cell isolation, this same region was excised and processed enzymatically (Figure 1C). Cell identity after isolation was also confirmed using immunostaining against HCN4. As shown in Figure 1D, besides HCN4-positive staining, pacemaker cells can be easily identified by their small and thin size compared to non-pacemaker cells. In addition, pacemaker cells exhibit three different morphologies: spindle, elongated, and spider, as other groups have reported^22,23^, which are easily distinguishable from large and square HCN4-negative cells (Figure 1E). Another useful criterion for identifying pacemaker cells is that they exhibit less prominent striations, which can be observed with bright-field illumination, and it is more evident when cells are stained against RyR2 (Figure 1D-E). For fixed-cell experiments, pacemaker identity was corroborated by HCN4 co-immunostaining; only HCN4-positive cells were included. For live-cell experiments, cells were selected based on their morphology and striation pattern.

### Aging slows down the firing rate and the diastolic depolarization of pacemaker cells

We recorded the spontaneous electrical activity of isolated pacemaker cells from 4-6 and 28-30 months old mice (Figure 2A). These ages are equivalent to 23-30 and 76-81 years in humans^24^. All recordings were acquired between 32-34°C. Figure 2B illustrates the parameters measured from the action potential waveforms to compare young and old cells. From all the analyzed parameters, significant changes in the old group were found for the action potential firing rate, the diastolic depolarization (DD) duration, the slopes of the Early (EDD) and Late (LDD) diastolic depolarization, the action potential duration (APD), and the repolarization duration (Figure 2C). At the population level, the action potential firing rate was significantly slower in old cells with a mean ± SEM value of 150 ± 20 bpm compared to 235 ± 16 bpm in young cells (Figure 2D). Firing rate from young animals was comparable to that reported by Marger et al (average of 260 bpm at 36°C)^25^. In addition to a slower firing rate, old cells showed an irregular rhythm evidenced by a higher standard deviation (SD) of the cycle length, with a value of 375 ± 127 ms for the old compared to 87 ± 14 ms for the young (Figure 2E). Since the DD duration and rate of depolarization (i.e., slope) are key determinants of how fast pacemaker cells can fire, we evaluated the differences between these parameters in young and old cells. Old cells showed a longer DD duration, with an average of 199 ± 24 ms compared to 106 ± 9 ms for the young (Figure 2F). We estimated the change in slope between the two age groups by fitting linear regression equations to the early and late DD. Figure 2G exemplifies the flattening of the early DD and late DD slopes observed in old pacemaker cells. We found that old cells have a significant reduction in the slopes of both Early DD (young 0.075 ± 0.007 mV/ms vs old 0.024 ± 0.004 mV/ms, Figure 2H) and Late DD (young 0.68 ± 0.08 mV/ms vs old 0.34 ± 0.03 mV/ms, Figure 2I). The action potential in old cells also lasted longer (young 144 ± 9 ms vs old 183 ± 19 ms).

**Figure 2.**
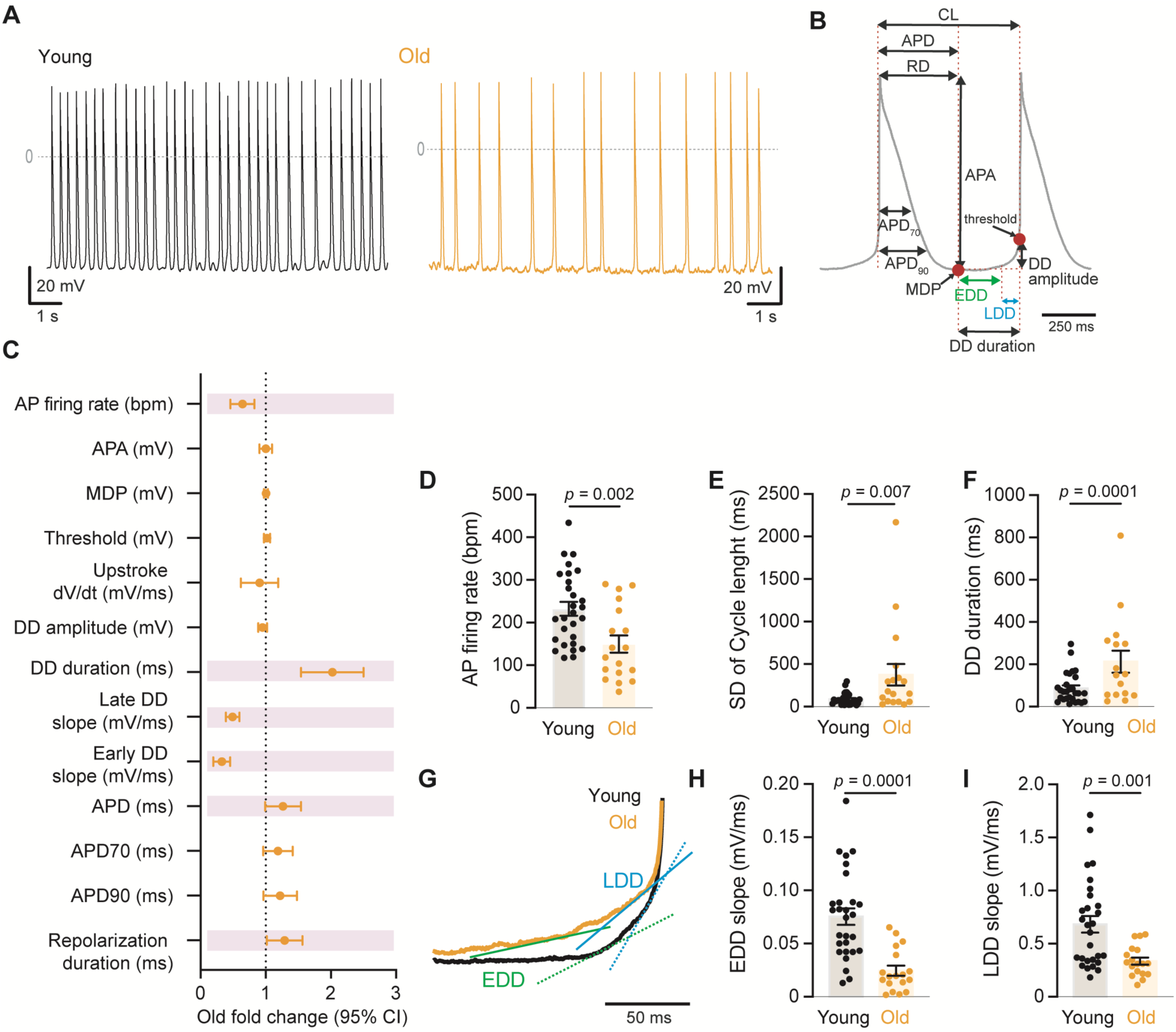
Aging slows down the action potential firing rate and the diastolic depolarization of pacemaker cells. **A.** Representative recordings of spontaneous action potential from young (black) and old (orange) pacemaker cells. **B.** Schematic of two pacemaker action potentials depicting the parameters analyzed (CL: cycle length, APD: action potential duration, RD: repolarization duration, APA: action potential amplitude, MDP: maximum diastolic depolarization, DD: diastolic depolarization, EDD: Early DD, LDD: Late DD. **C.** Fold change of the action potential parameters analyzed in old pacemaker cells relative to young. The vertical dotted line represents the young population. Purple bars highlight parameters that were significantly different from the young. **D.** Comparison of the firing rate from young and old pacemaker cells. **E**. Comparison of the standard deviation (SD) of the cycle length from young and old pacemaker cells. **F.** Comparison of the DD duration from young and old pacemaker cells. **G.** Representative traces of the diastolic depolarization phase from a young and old pacemaker cell to compare the Early DD (green) and Late DD (blue) slopes. Adjusted lines were used to calculate the depolarization rate of each phase. **H, I.** Comparison of the Early DD and Late DD slopes from young and old pacemaker cells. In all the scattered plots, bars represent the mean, and error bars the SEM. Statistical comparisons used a two-tailed t-test comparing a population of n = 28 cells, N = 5 mice for the young group and n = 18 cells, N = 5 mice for the old.

### L-type calcium current density is reduced in old pacemaker cells

The flattening of the DD observed in old pacemaker cells suggests that aging affects the ion channels responsible for this spontaneous depolarization. Three types of ion channels play a significant role in this phase: HCN4, T-type, and L-type calcium channels. The effect of aging on HCN4 channels is well known. Given that mutations in the L-type Ca_V_1.3 channel lead to bradycardia, similar to what is observed in the elderly^14,15^, we focused on the effect of aging on L-type calcium channels. Our specific aim was to determine whether isolated pacemaker cells from young and old animals differed in their calcium current density. Whole-cell calcium currents were recorded from isolated pacemaker cells in response to 20 ms depolarizing pulses to -20 mV from a holding potential of -80 mV. Figure 3A shows representative traces of the total calcium current (I_Ca TOTAL_) in young and old pacemaker cells.

**Figure 3.**
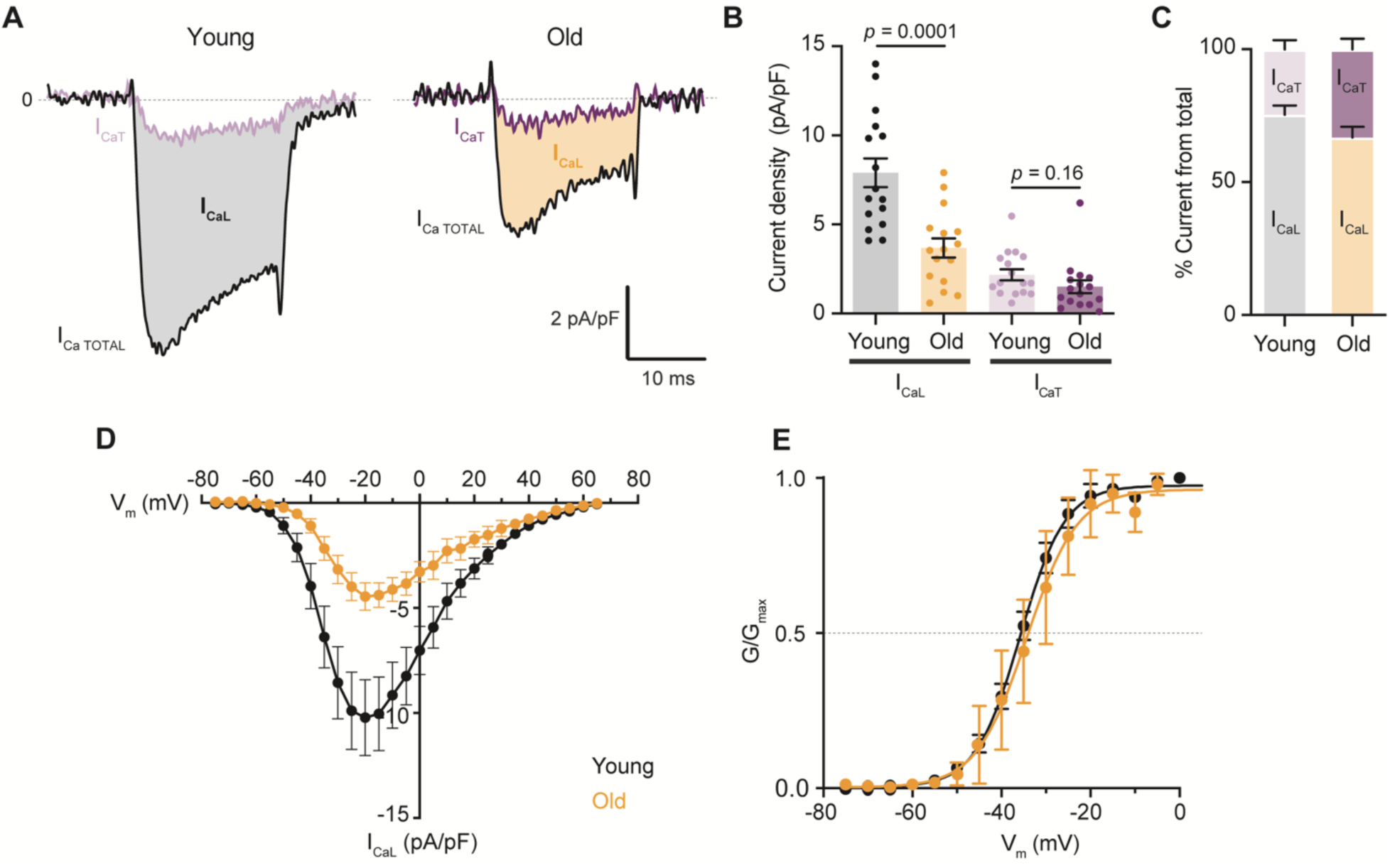
L-type calcium current density is reduced in old pacemaker cells. **A.** Representative recordings of the total calcium current (I_Ca TOTAL_, black traces) and the nifedipine-resistant component (I_CaT_, light and dark purple traces) from young and old pacemaker cells. The shaded region of the current corresponds to the nifedipine-sensitive component (I_CaL_, grey for young and orange for old). **B.** Comparison of calcium current density with age for both L- and T-type. **C.** Percentages of I_CaL_ and I_CaT_ relative to total calcium currents observed in each age group. Note that in this plot, total calcium current in old cells is also shown as 100%, but it is only 50% of the current observed in young cells. **D.** Comparison of the current-voltage relationship for I_CaL_ from young and old pacemaker cells. **E.** Normalized conductance (G) - voltage relationship for I_CaL_ from young and old pacemaker cells. In all the scattered plots, bars represent the mean, and error bars the SEM. Statistical comparisons used a two-tailed t-test comparing a population of n = 16 cells, N = 4 mice for the young group and n = 16 cells, N = 4 mice for the old.

To isolate the L-type calcium component, calcium currents were blocked by bathing the cells with an external solution containing 10 μM of the specific L-type blocker nifedipine. Nifedipine has been shown to block some T-type channels. However, the Ca_V_3.1 isoform of T-type calcium channels, α1G, which is the predominant isoform in sinoatrial node pacemaker cells, is blocked with an IC_50_ of 109 μM, which is 10-fold more than the concentration we used in these experiments. Furthermore, A high concentration of nifedipine (100 μM) blocks Ca_V_3.1 only by 23%. For these reasons, the conditions of our experiments are not expected to block the T-type component.

As shown in the light and dark purple traces in Figure 3A, only a small component of the I_Ca TOTAL_ is resistant to nifedipine and, hence, carried by T-type calcium channels (I_CaT_). The L-type component of the current (I_CaL_) was calculated by subtracting the nifedipine-resistant component from the I_Ca TOTAL_. The analysis of the I_CaL_ and I_CaT_ current densities showed that aging reduced the I_CaL_ by 53%, going from 7.9 ± 0.8 pA/pF in young cells to 3.7 ± 0.5 pA/pF in the old (Figure 3B). In contrast, I_CaT_ was not significantly different, with values of 2.2 ± 0.3 pA/pF in young cells and 1.5 ± 0.4 pA/pF in old. The percentage of total calcium current carried by each component is shifted with age, with a larger contribution of T-type in old cells (Figure 3C). We also evaluated the effect of aging on the voltage dependence of the L-type calcium current. For this, we constructed I-V and G-V L-type calcium currents curves in response to increasing 5 mV depolarizing pulses from a holding potential of -75 mV to +65 mV. Old pacemaker cells had a significant reduction of the L-type calcium current at all voltages with no shift of the voltage dependence (Figures 3D, 3E). We next sought to discover the underlying mechanism behind the reduction in the L-type calcium current.

### Aging does not lead to cell hypertrophy

The effects of aging on the size of the heart are widely recognized. A well-accepted age-associated alteration is increased ventricular wall thickness^26–28^, which is underlined by cardiomyocyte hypertrophy. Age-associated structural changes in the pacemaker have been previously reported, including loss of cell number^29^ and an increase in fibrosis^30^. Cell hypertrophy in old pacemaker cells could explain the observed reduction in L-type calcium current density. We sought to test this hypothesis and determine the effects of age on pacemaker cell size.

To assess changes in cell size in the pacemaker, we used three different strategies. First, we quantified the cell size from high-resolution volumetric reconstructions of the sinoatrial node explant. Explants were isolated and immunostained against HCN4 (Figure 4A). This signal is detected in the cell periphery as expected from a protein localized to the plasma membrane. Individual cells had an average width of 6.2 ± 0.3 μm in young and 6.7 ± 0.3 μm in old mice (Figure 4B), and an average length of 103 ± 3 μm in young and 112 ± 5 μm in old mice (Figure 4C). These morphological parameters were not significantly different. As a second approach, we measured the same morphological parameters from single, isolated pacemaker cells immunostained after tissue dissociation (Figure 4D-G). We found that neither the width (7.1 ± 0.3 μm in young vs 8.4 ± 0.3 μm in old), length (96 ± 4 μm in young vs 99 ± 3 μm in old), nor the area (627 ± 36 μm^2^ in young vs 702 ± 33 μm^2^ in old) was significantly different in isolated cells. As a last approach to evaluate changes in cell size, we quantified the cell capacitance, an electrophysiological measurement directly proportional to cell area. The capacitance was non-significantly different between young (26.8 ± 2.0 pF, n = 22) and old (27.2 ± 1.8 pF, n = 21) pacemaker cells (Figure 4H). These results show that pacemaker cells do not undergo cellular hypertrophy and therefore, the reduction in L-type calcium current is not due to this mechanism.

**Figure 4.**
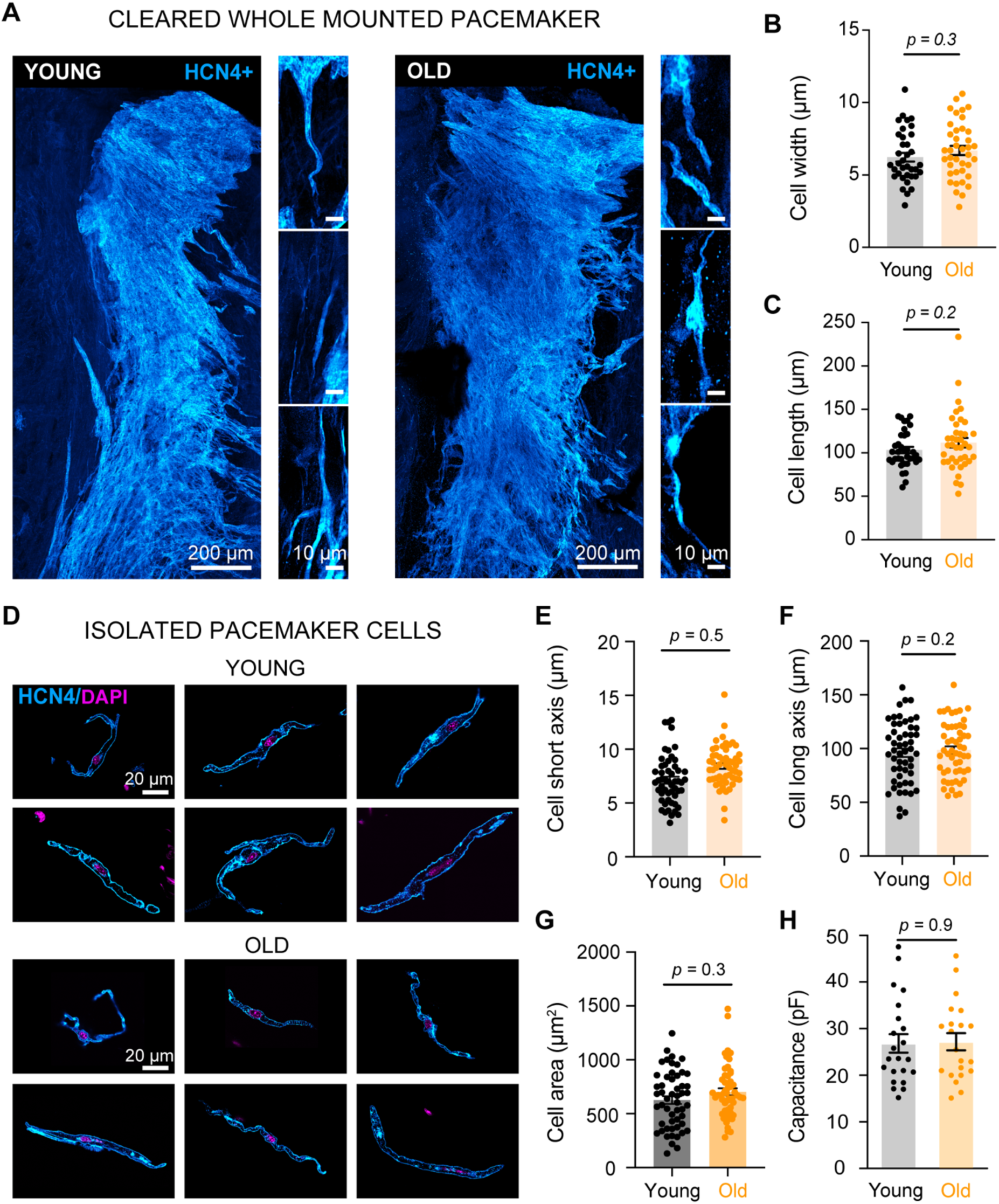
Aging does not lead to cell hypertrophy. **A.** Pictures of heart explants showing the pacemaker region stained against HCN4 in young (left) and old (right) mice. **B.** Comparison of cell width from HCN4-positive cells measured from tissue explants. **C.** Comparison of cell length from HCN4-positive cells measured from tissue explants. Measurements in B - D were performed from three tissues in each age, and the total number of cells was 37 in the young and 39 in the old. **D.** Pictures of isolated pacemaker cells from young (top) or old (bottom) mice stained against HCN4. Comparison of short axis (**E**), long axis (**F**), and cell area (**G**) from isolated HCN4-positive cells. Measurements in panels E-G are from 53 pacemaker cells in young and 54 pacemaker cells in old. **H.** Comparison of the capacitance of young and old pacemaker cells. Sample numbers were n = 22 cells, N = 4 mice for the young group, and n = 21 cells, N = 4 mice for the old.

### Reduction in L-type calcium current in old pacemaker cells is associated with a decrease in their density and clustering at the plasma membrane

We evaluated whether the observed age-associated reduction in current density was caused by a reduction in the expression of L-type calcium channels. We measured the Ca_V_1.2 and Ca_V_1.3 protein levels by western blot from protein lysates obtained from sinoatrial node explants from young and old animals. Contrary to our prediction, total protein levels were not decreased in samples from old animals. On the contrary, Ca_V_1.2 channels were 1.9 ± 0.4-fold higher in old animals compared to the young. The expression of Ca_V_1.3 channels was the same in old and young animals (0.9 ± 0.2-fold relative to the young, Figures 5A and 5B).

**Figure 5.**
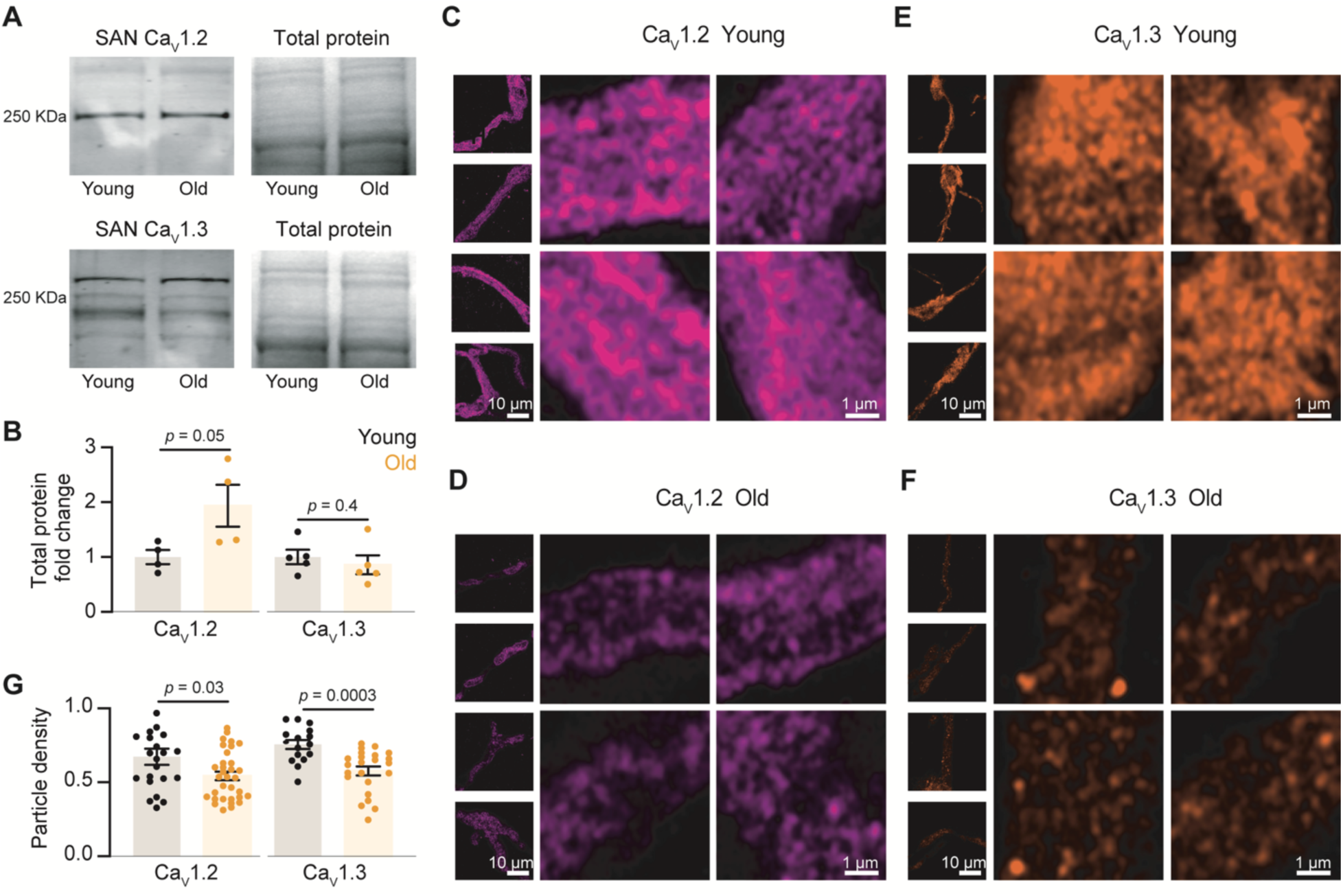
Aging does not affect the total expression of L-type calcium channels but reduces their density near the plasma membrane. **A.** Representative western blots for Ca_V_1.2 and Ca_V_1.3 channels and total protein stains from pacemaker tissue explants from young and old mice. **B.** Fold change of Ca_V_1.2 and Ca_V_1.3 channel total expression in old animals relative to young. Expression was normalized to total protein and each data point represents an animal. Statistical comparisons used a two-tail Mann-Whitney test. **C-F.** Representative AiryScan high-resolution images of the footprint of young and old pacemaker cells labeled against Ca_V_1.2 (magenta) or Ca_V_1.3 (orange). Insets are 5 μm by 5 μm footprint regions from each of the cells shown to the left. **G.** Comparison of Ca_V_1.2 and Ca_V_1.3 particle density between young (Ca_V_1.2, n = 22, N = 3; Ca_V_1.3, n = 16, N = 3) and old (Ca_V_1.2, n= 33, N = 3; Ca_V_1.3, n = 23, N = 3) cells. Statistical comparisons used a two-tail t-test.

As the total expression of the channels did not seem to be altered, we next assessed whether channel expression at the plasma membrane was reduced with aging. As a first approach, we used high-resolution Airyscan microscopy to compare the fluorescence density at the cell footprint in cells immunostained with antibodies against Ca_V_1.2 or Ca_V_1.3 channels. The labeling for calcium channels was not homogeneous; it exhibited a dotted distribution. We quantified and compared the particle density and observed that old cells showed 19% fewer Ca_V_1.2 particles per unit of membrane area, with a density of 0.54 ± 0.03 particles/μm^2^ compared to 0.67 ± 0.05 particles/μm^2^ in the young (Figures 5C, 3D, and 5G). Ca_V_1.3 particle density was reduced in old pacemaker cells by 24%, with values of 0.75 ± 0.03 particles/μm^2^ for the young and 0.58 ± 0.03 particles/μm^2^ for the old (Figures 5E, 5F, and 5G).

As a second approach, we compared the organization of L-type calcium channels at a nanometer resolution. Our group previously showed that L-type Ca_V_1.2 and Ca_V_1.3 channels organize in packed clusters at the plasma membrane in several cell types, including ventricular cardiomyocytes and hippocampal neurons^31–34^. Using Stochastic Optical Reconstruction Microscopy in Total Internal Reflection Mode, we quantified the density of calcium channels with 20 nm lateral resolution and 50 nm axial resolution in an area restricted to the plasma membrane. We found that L-type calcium channels were organized in clusters at the plasma membrane of young pacemaker cells with an average size of 4265 ± 531 nm^2^ for the Ca_V_1.2 and 3975 ± 334 nm^2^ for the Ca_V_1.3 (Figures 6A-C). Cells from old animals showed a significant reduction in the mean cluster size with values of 2416 ± 342 nm^2^ for Ca_V_1.2, and 2321 ± 132 nm^2^ for Ca_V_1.3 channels (Figures 6A-C). The number of channels per membrane unit area was also reduced in old cells. For the Ca_V_1.2 channels, cluster density was reduced by 38%, going from 9.6 ± 1.7 clusters/μm^2^ in young cells to 5.9 ± 0.5 clusters/μm^2^ in the old (Figure 4D). For Ca_V_1.3 channels, cluster density was reduced by 39%, going from 9.7 ± 0.9 clusters/μm^2^ in young to 5.9 ± 1.3 clusters/μm^2^ in old cells (Figure 6D). Figures 6E and 6F show the frequency distribution for the Ca_V_1.2 and Ca_V_1.3 channel cluster area. The cluster size frequency distributions were, in both cases, shifted down and to the left in old cells, suggesting an age-associated global reduction in the L-type channel cluster size. Taking together the results from these two approaches, we concluded that aging reduces the number of L-type calcium channels inserted at the plasma membrane but not their total expression, and that this reduction in density at the plasma membrane leads to a reduction in the number of channels found in clusters.

**Figure 6.**
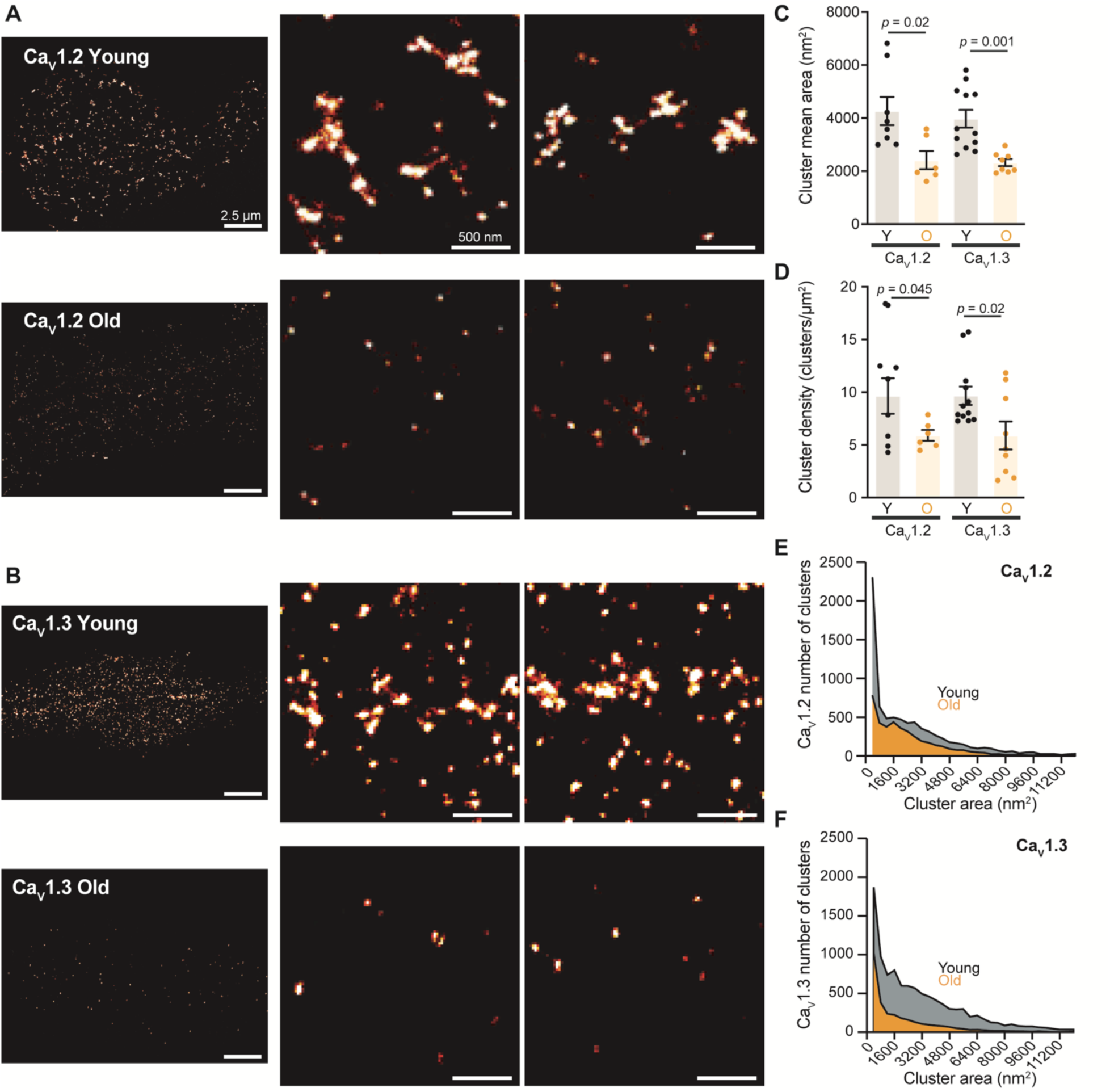
Aging reduces L-type calcium channel clustering. **A** and **B.** Representative super-resolution images of Ca_V_1.2 and Ca_V_1.3 channels in young and old pacemaker cells. Panels to the right of each image are zoom in. **C.** Average cluster area for Ca_V_1.2 and Ca_V_1.3 channel clusters in young and old pacemaker cells. **D.** Comparison of Ca_V_1.2 and Ca_V_1.3 cluster density between young and old cells. **E and F.** Comparison of the frequency distributions of the area of Ca_V_1.2 and Ca_V_1.3 channel clusters between young and old cells. In all the scattered plots bars represent the mean and error bars the SEM. Statistical comparisons used a two-tail t-test comparing a population of n = 8 cells, N = 3 mice for Ca_V_1.2 young; n = 12 cells, N = 4 mice for Ca_V_1.3 young; n = 6 cells, N = 3 mice for Ca_V_1.2 old; and n = 8 cells, N = 3 mice for Ca_V_1.3 old.

### Reduction in L-type calcium current in old pacemaker cells is associated with a decrease in the activity of single channels

To further test the hypothesis that aging leads to a reduction of channels at the plasma membrane and that fewer channels result in a reduction in L-type calcium current, we assessed the activity of single channels in small membrane patches. This strategy provides a functional readout of the number of channels found in that membrane area and their probability of opening. Figure 7A shows representative single-channel recordings from young and old pacemaker cells. The channel activity in patches from young cells was characterized by coordinated opening of multiple channels. Notably, single-channel recordings in old cells were dominated by independent single-channel events that rarely exhibited coordinated opening of multiple channels. To compare the activity between the age groups, we calculated the parameter NP_o_ (the number of channels times the open probability) for each patch. This parameter was around 4 times larger in young (0.23 ± 0.06) than old (0.06 ± 0.01) patches (Figure 7B). The change in NP_o_ measured by patch electrophysiology was considerably larger than the change in channel density measured by microscopy, suggesting that aging not only affects the number of channels but also their open probability.

**Figure 7.**
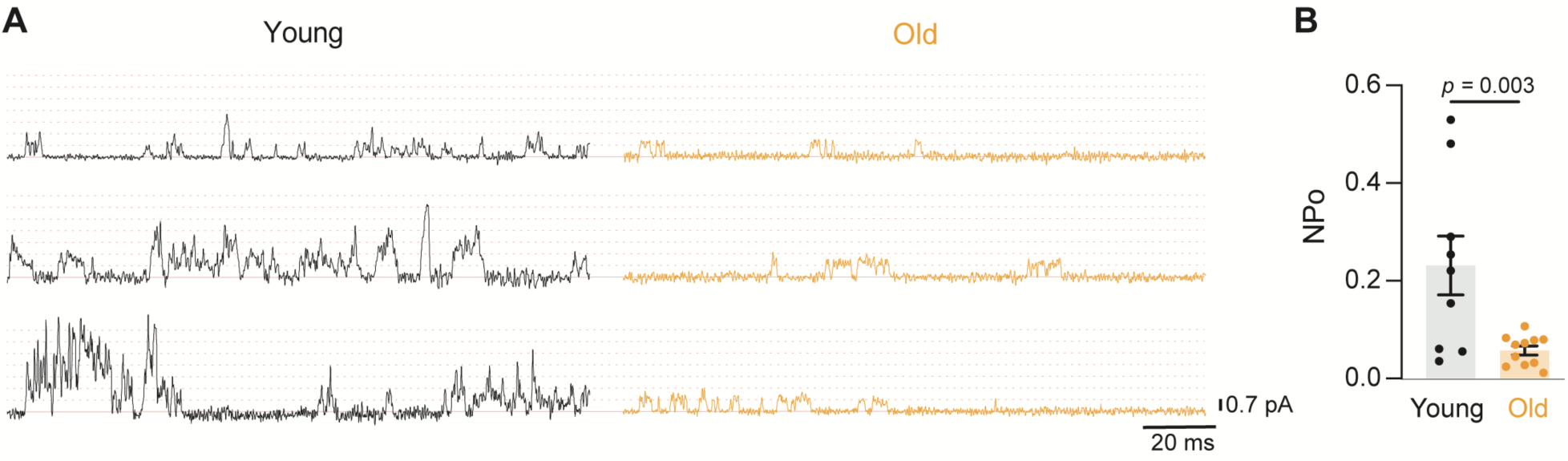
Aging reduces the number of active channels at the plasma membrane and their open probability. **A.** Representative single-channel recordings from young (black) and old (orange) pacemaker cells. Dotted lines represent number of channels opening (1-7 channels), from the baseline closed state based on a unitary current of 0.7 pA. **B.** Comparison of the channel open probability (NP_Ο_) between young and old pacemaker cells calculated from 50 voltage sweeps per cell. Statistical comparisons used a two-tail t-test comparing a population of n = 9 cells, N = 3 mice for the young group and n = 8 cells, N = 3 mice for the old group.

### Increasing L-type calcium channel open probability restores firing rates in pacemaker cells from old animals

If a decrease in the open probability of L-type calcium channels is one of the factors by which aging slows down the pacemaker rate, one would predict that increasing it in old pacemaker cells would accelerate their firing rate. To test this hypothesis, we recorded the spontaneous firing rate of isolated young and old pacemaker cells in control conditions and under the bath application of 500 nM of the L-type channel agonist Bay K 8644. Bay K has been shown to increase the open probability and opening time of L-type calcium channels^35–40^. In this new experimental set under control conditions, we found that the rate was slower in cells from old animals compared to young, as we had shown before (Figure 8A, left). The perfusion of Bay K accelerated the spontaneous firing rate in young cells 1.3-fold, going from 209 ± 22 bpm to 283 ± 20 bpm (Figure 8B). The effect of Bay K in old cells was stronger, with an acceleration of 1.6-fold, going from 151 ± 27 to 242 ± 19 bpm (Figures 8A and 8B). Remarkably, increasing L-type calcium channel open probability with Bay K was enough to accelerate the firing rate in old cells to values not significantly different from the young group (Figure 8B). Increasing the open probability of L-type calcium channels had a similar effect on other action potential parameters. The application of Bay K reverted the significant differences observed in control conditions between young and old for the DD duration, Early DD and Late DD slopes, and repolarization duration (Figure 8C). Given the slowdown of the Early DD and Late DD in old cells shown in Figure 2, we analyzed the effect of Bay K on the DD duration and the slopes of the Early and Late DD. Bay K changed the kinetics of Early DD and Late DD in old cells, making them similar to those observed in the young (Figure 8D). A paired analysis of the DD showed that Bay K significantly shortened DD only in old cells (Figure 8E). Bay K induced a significant increase in the Late DD and Early DD slopes in both groups (Figures 8F and 8G). In control conditions, the Early DD and Late DD slopes were significantly less steep in old cells when compared to the young (Early DD: 0.02 ± 0.1 mV/ms in old vs 0.07 ± 0.01 mV/ms in young; Late DD: 0.30 ± 0.04 mV/ms in old vs 0.60 ± 0.08 mV/ms in young) (Figures 8F and 8G). However, under Bay K application, Early DD and Late DD were significantly accelerated in the old group, reverting the significant differences relative to the young (Early DD: 0.09 ± 0.01 mV/ms in young vs 0.06 ± 0.01 mV/ms in old; Late DD: 0.74 ± 0.09 mV/ms in young vs 0.58 ± 0.01 mV/ms in old cells and). Early DD and Late DD were accelerated 1.4 and 1.2 times, respectively, in young cells. In contrast, Bay K accelerated Early DD 2.6 times and Late DD 1.9 times in old cells. The differences in the repolarization time were also reversed upon Bay K treatment (Figure 8H). Together, these results suggest that increasing the open probability of L-type calcium channels in old pacemaker cells contributes to pacemaker rate acceleration through an increase in the diastolic depolarization rate combined with a reduction in the time for repolarization.

**Figure 8.**
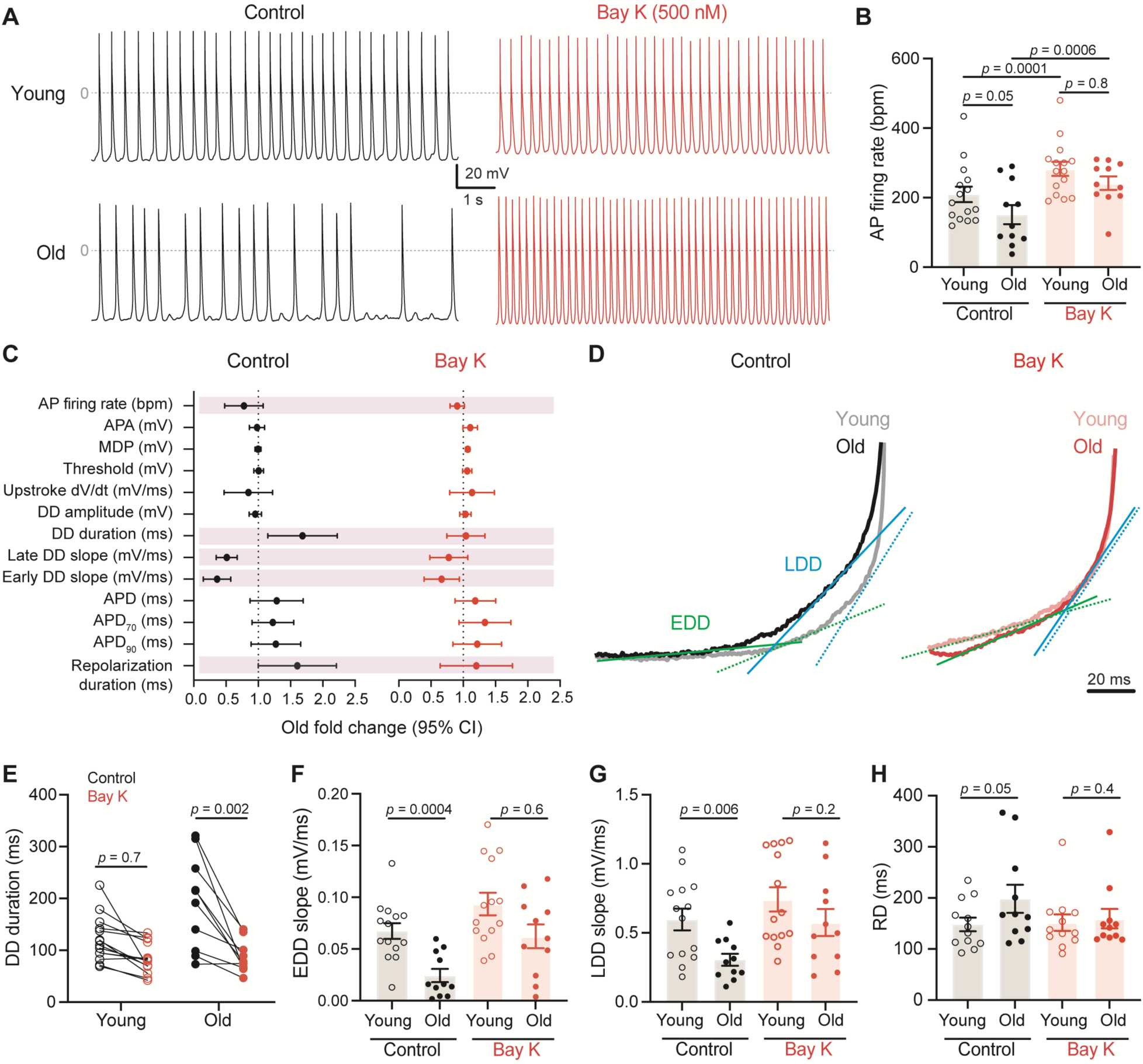
Increasing L-type calcium channel open probability restores normal pacemaking rates in old cells. **A.** Representative recordings of action potentials from young and old pacemaker cells before (black) and after (red) application of 500 nM Bay K8644. **B.** Comparison of the effect of Bay K on the firing rate of young and old pacemaker cells. **C.** Fold change of the action potential parameters analyzed in old pacemaker cells relative to young in control (black) and Bay K (red). Vertical dotted lines represent the young population. Purple bars highlight parameters that were significantly different in control conditions but reverted by Bay K. **D.** Representative overlapped magnification of the diastolic depolarization in the absence (control) and presence of Bay K. Straight lines illustrate how the kinetics of the Early DD (green) and Late DD (blue) phases were calculated from young (dotted) and old (solid) traces. **E.** Comparison of the effect of Bay K (red) on the duration of the diastolic depolarization (DD) relative to control (black) in young and old pacemaker cells. **F,G**. Comparison of the slopes of the tangential lines for Early DD (F) and Late DD (G) between young and old pacemaker cells in control conditions (black) and upon the subsequent addition of Bay K (red). **H.** Comparison of the repolarization duration (RD) between young and old pacemaker cells in control conditions (black) and upon the subsequent addition of Bay K (red). In all the scattered plots bars represent the mean and error bars the SEM. Statistical comparisons between young and old used a two-tail t-test comparing a population of n = 14 cells, N = 3 for the young and n = 11 cells, N = 3 mice for the old. Statistical comparisons between control and Bay K used a paired two-tail t-test.

Together, these results suggest that a reduction in the open probability of L-type calcium channels is one of the mechanisms by which aging slows down the firing frequency of pacemaker cells.

## Discussion

Our work centered on understanding the molecular mechanisms behind the slowdown of the cardiac pacemaker. We have contributed to answering this question by providing additional evidence that the L-type calcium current is reduced with aging. We also provide evidence to understand the mechanism behind this reduction. The age-associated mechanism is twofold: a reduction of the number of channels at the plasma membrane and an impairment of their function, more specifically, their open probability. Although we could not unveil the mechanisms behind the observed change in number and activity of these channels, our results allowed us to rule out some potential explanations and suggest a path forward. Our results rule out cell size, total channel expression, and voltage-dependence alterations. Instead, we observed that the low expression of channels at the plasma membrane reduces their clustering.

Our results show that cardiac pacemaker cells do not undergo hypertrophy. This conclusion disagrees with two previous publications. The discrepancy may be due to differences in species, age under study, or the techniques used. We compared two age groups, 4-6 months and 28-32 months, of C57BL/6 mice. In one study, the cell size was found to be 33% larger in 28-month-old mice compared to 2-month-old mice^1^. Measurements were taken with electrophysiological equipment and reported as cell capacitance. This is considered a highly sensitive technique, as it can detect minute changes in cell capacitance (cell size) as little as the addition of membrane from a single vesicle, which would be impossible to detect using microscopy. One limitation of this approach is that the cell size and shape are expected to change as the recording solution from the pipette dialyzes into the cell. A different approach was used to study the effect of aging on cell morphology in rats. In this study, it was found that old pacemaker cells are 38% larger than young ones^41^. This study compared three and 24-month-old rats. Size was determined by measuring cell diameter in cross sections of pacemaker tissue stained against caveolin-3 antibody or Masson’s trichrome staining. What then makes us confident of our results? Given the discrepancy, we used three complementary approaches, and the results were consistent. We used a modern microscopy technique to measure cell size in situ, leaving the tissue intact. Instead of sectioning the tissue, we stained against HCN4 and cleared the tissue. This technique allowed us to measure cell length and width. Moreover, we imaged the tissue using a high-resolution microscope, improving the effective resolution. This technical improvement should allow for the detection of even smaller changes in size. In addition to measuring cell size in the tissue, we confirmed the results by measuring length, width, and area in isolated cells. In isolated cells, we imaged the cells using a higher magnification objective. This approach provided better axial resolution and a large sample size; the results in isolated cells were consistent with those obtained in the tissue, ruling out possible effects on cell size caused by the isolation protocol. Finally, we also compared the capacitance of young and old cells and found consistent results.

If aging affects the channels directly, and not through changes in cell size, what are the mechanisms involved? Our results indicate that aging causes a decrease in the channels at the plasma membrane. This reduction could be explained by a reduction in the insertion or an increase in the removal of channels. One of the hallmarks of aging is the loss of proteostasis^42,43^. Misfolding or errors in translation of the channels could cause degradation or problems in trafficking of the proteins, preventing them from being inserted correctly at the plasma membrane. This loss of proteostasis in the heart is strongly associated with cardiac dysfunction^44^. Furthermore, the proteins of the aging heart exhibit increased oxidation^44,45^ and ubiquitation^46^, common aspects of the loss of proteostasis in aging. Moreover, the proteome of the heart is remodeled in aging. While proteins of the electron transport chain, citric acid cycle, and fatty acid metabolism are decreased, proteins in glycolysis and extracellular structural proteins are increased^47^. These changes point to an alteration in protein quality control systems, leading to accumulation of damaged proteins. Our results, then, open a new scenario in which the effects of aging might alter ionic currents not only through changes in expression, but through deficits of folding, traffic, and degradation.

The observed reduction in the number of calcium channels at the plasma membrane and their organization in clusters could also be explained by a disruption in specialized domains that influence ion channel insertion and organization. One of these special regions is formed by caveolin proteins. In contrast to working cardiomyocytes, pacemaker cells lack T-tubules. However, they have an extensive caveolar network that serves as specialized signaling microdomains where many channels, including L-type calcium channels, reside. Aging has been linked to the disruption of this caveolar network and its effects in the heart and other tissues^48–51^. In ventricular cardiomyocytes, the loss of caveolin-3 has been linked to the age-associated reduction of calcium current in the T-tubules^52^. As we have previously shown, aging causes a reduction in the caveolar network in pacemaker cells, accompanied by a decrease in the interaction between caveolin-3 and the calcium channels Ca_V_1.2 and Ca_V_1.3^53^. The role of an age-associated loss of caveolin-3 in the insertion and removal of ion channels expressed in pacemaker cells needs further investigation.

As observed by super-resolution microscopy, another potential factor contributing to smaller calcium channel clusters involves the alteration of scaffolding proteins. Scaffold proteins identified to bind L-type calcium channels in other cell types include BIN1^54,55^, AKAPs^56–60^, and Shank^61–63^. BIN1 plays an important role in the delivery and clustering of Ca_V_1.2 channels in ventricular cardiomyocytes, while AKAP150 has been shown to be necessary for the cooperative gating of Ca_V_1.2 channels in smooth muscle cells. However, the function of the majority of these scaffolds remains to be tested in pacemaker cells.

Our results show that L-type calcium channels exhibit a reduced open probability in old pacemaker cells. Notably, increasing the open probability with Bay K restored firing frequency in old cells to young levels, supporting our conclusion that the reduction in the open probability of L-type calcium channels is an important factor associated with the age-associated slowdown. What factors could be reducing the open probability of these channels? L-type calcium channels exist in two different modes^37^. Mode 1 is characterized by short-lasting openings (low Po), while mode 2 is characterized by long-lasting openings (High Po). Interestingly, the calcium channel agonist Bay K increases the frequency of finding the channels in mode 2 in both single-channel recordings and calcium sparklets^64^. This high activity mode represents the activation of channels that are organized in clusters. In the context of our results, we speculate that aging is affecting the open probability via a decrease of Mode 2. We have previously shown that both Ca_V_1.2 and Ca_V_1.3 channels share a facilitation mechanism that depends on the formation of these clusters^31,32^. Proximity between L-type calcium channels allows for a physical interaction between their C-terminal domains mediated by calmodulin. This functional coupling directly increases the open probability of the channels, increasing the influx of calcium into the cell. Therefore, the age-associated reduction of the open probability could be caused by decreased functional coupling between nearby channels. As the functional coupling depends on the availability of adjacent channels to interact, our results support the hypothesis that a reduction in cluster size can lead to a decrease in the total number of coupled calcium channels. However, careful characterization of the channel coupling in old pacemaker cells is needed to test this idea further.

## Methods and Materials

### Solutions

For the isolation of pacemaker cells, the following solutions were used: Tyrode’s III solution (in mM) – 148 NaCl, 5.4 KCl, 5 HEPES, 5.5 glucose, 1 MgCl_2_, and 1.8 CaCl_2_, pH was adjusted to 7.4 with NaOH. Low-Calcium Tyrode’s solution (in mM) – 140 NaCl, 5.4 KCl, 5 HEPES, 5.5 glucose, 1.2 KH_2_PO_4_, 50 taurine, 0.5 MgCl_2_, and 0.2 CaCl_2_, pH was adjusted to pH 6.9 with NaOH. KB solution (in mM) – 80 L-glutamic acid, 25 KCl, 10 HEPES, 10 glucose, 10 KH_2_PO_4_, 20 taurine, 0.5 EGTA, and 3 MgCl_2_, pH was adjusted to 7.4 with KOH.

For electrophysiological recordings, the following solutions were used: Whole-cell voltage-clamp calcium current (I_Ca_) external solution (in mM) – 110 N-methyl-D-glutamine, 5 CsCl, 10 HEPES, 10 glucose, 30 TEA-Cl, 4 4-Aminopyridine, 1 MgCl_2_, and 2 CaCl_2_, pH was adjusted to 7.4 with HCl. I_Ca_ internal solution (in mM) – 50 CsCl, 10 HEPES, 70 L-Aspartic acid, 30 TEA-Cl, 5 EGTA, 5 Mg-ATP, 1 MgCl_2_, and 0.7 CaCl_2_, pH was adjusted to 7.2 with CsOH. Perforated-patch current-clamp action potential external solution was Tyrode’s III. Perforated-patch current-clamp action potential internal solution (in mM) – 130 L-Aspartic acid K, 10 NaCl, 10 HEPES, 0.04 CaCl_2_, 2 Mg-ATP, 0.1 Na-GTP, 6.6 phosphocreatine, pH was adjusted to 7.2 with KOH. Cell-attached single channel recording external solution (in mM) – 145 KCl, 10 NaCl, 10 HEPES, pH was adjusted to 7.4 with NaOH. Cell-attached single channel recording internal solution (in mM) – 110 CaCl_2_, 20 TEA-Cl, 10 HEPES, and 10 μM of the HCN channel-blocker ivabradine, pH was adjusted to 7.2 with CsOH.

For protein extraction, the following solutions were used: Homogenization buffer (in mM) – 150 NaCl, 50 Trizma base (from 1M stock at pH 7.8 with HCl), 5 EDTA (from 50 mM stock pH 7.8 with NaOH); the day before the experiment NaF 10 mM, Glycerol 5% v/v, Triton X-100 1% v/v, Sodium deoxycholate 0.25% were added; on the day of the experiment one tablet of SIGMAFAST^TM^ protease inhibitor cocktail (Sigma, S8830), EDTA free in 50 ml of the buffer was added. Lysis Buffer (to make 10 ml) – 10 ml of homogenization buffer, 8.3 μl Calpain I (10 mg/ml) (Fisher, AAJ61766LB0), 8.3 μl Calpain II (10 mg/ml) (Sigma, A6060), 10 μl o-phenathroline (200 mg/ml) (Sigma, 33510), 10 μl PMSF (20 mg/ml) (Sigma, P7626), 10 μl Benzamidine (15.7 mg/ml) (Sigma, 12072), prepared the day of the experiment. Solutions were kept on ice.

### Pacemaker cells isolation

Pacemaker cells were freshly isolated from mature adults (4-6 months old), or old (24-28 months old) C57BL/6 male mice. All work with animals was performed under an approved protocol by the University of Washington Institutional Animal Care and Use Committee (IACUC). Animals were euthanized by intraperitoneal overdose of Euthasol (Virbac, 400mg/ml). The heart was dissected and pinned down on a Sylgard-coated dish containing warm Tyrode’s III solution. The sinoatrial node region was identified under the stereoscope as the region delimited at the top by the superior vena cava, at the right by the sulcus terminalis, and the bottom by the coronary sulcus and the inferior vena cava (Figure 1A). The sinoatrial node artery was also used as an anatomical landmark to identify the pacemaker region. Pacemaker cells were then isolated from the sinoatrial node following the protocol described by Fenske et al. 2016^65^. Briefly, the excised sinoatrial node tissue was immersed in a 37°C pre-heated 2 ml tube containing 675 μl of Low-Calcium Tyrode’s. After 5 min of stabilization at 37°C the following compounds were added to the tube: BSA (Cf = 1 mg/ml), elastase (Cf = 18.87 U/ml, Millipore 324682), protease (Cf = 1.79 U/ml, Sigma P5147), and collagenase B (Cf = 0.54 U/ml, Roche 11088807001). The enzymatic digestion of the tissue was carried out for 30-35 minutes in a 37°C water bath, and a mechanical dissociation with a short fire-polished glass pipette was performed every 7 minutes during incubation. To stop the digestion, the digested tissue was centrifuged at 200 × g for 2 min at 4°C, and the supernatant was discarded and replaced with 1 ml of Tyrode’s Low-Ca^2+^ solution; this process was repeated twice. Then the tissue was washed two times more with 1 ml of Calcium-free KB solution. The dissociated tissue was left to rest in KB solution at 4°C for a minimum of 40 min. Finally, single pacemaker cells were resuspended by applying gentle mechanical dissociation with a flame-forged glass pipette. For electrophysiology experiments, to recover the automaticity of the pacemaker cells, calcium was reintroduced into the KB cell’s storage solution by the gradual addition of small amounts of Tyrode’s III solution (10, 50 and 100 μl at 5 min intervals). Cells were plated on poly-L-Lysine (PLL)-coated coverslips. PLL coating was performed at least one day before the cell isolation. Clean coverslips were incubated in PLL hydrobromide solution MW 300,000 (Sigma, P1524) for 30 min at 37°C. Each coverslip was then removed onto a rack and flushed with Milli-Q water and aspirated 5 times. Coverslips were left to dry in a biosafety cabinet overnight.

### Protein extraction and western blot

Animals were anesthetized, and the heart was quickly removed and placed in ice-cold homogenization buffer. The heart was pinned on a Sylgard-coated 60 mm dish containing ice-cold homogenization buffer. The sinoatrial node explant was dissected and quickly transferred into a glass micro-tissue grinder (DWK, 357844) containing 100 μl of lysis buffer. The tissue was homogenized on ice, and the sample was transferred to a 1.5 ml tube and placed in a rotator wheel at 4°C for two hours. Samples were centrifuged at 16,000 × g for 20 min at 4°C. The solubilized protein supernatant was transferred into a new tube and stored at -80°C. Protein concentration was determined by measuring 590 nm absorbance using Quick Start Bradford (Biorad Cat 5000205) in a Spectronic Genesis 5 plate spectrophotometer. Protein concentration obtained from a single sinoatrial node tissue ranged between 1.5 and 2.5 μg/μl. For electrophoresis, protein samples were denatured in SDS loading buffer for 10 min at 70°C. 25 μg of protein were loaded per line on a Mini Protean TGX Precast 4-10% Biorad gel. As a reference, the ladder spectra multicolor was loaded (Thermo, 26634). Gels were run for 15 min at 50 V, followed by 1 h at 130 V in TGS. Proteins were transferred onto 0.2 μm PVDF membranes (Thermo, 1704157) using an OwlTM Electroblotting system (Thermo, VEP-2) at 0.4 A and 4°C for 2 h. Membranes were stained for total protein using No-StainTM protein labeling reagent (Invitrogen, A44449), following the manufacturer’s instructions. Total protein images were taken using an iBright imaging system (Thermo). For immunoblotting, membranes were blocked in 5% Milk in TBST at RT for 1h with constant agitation. Rabbit polyclonal anti-Ca_V_1.2 (anti-CNC1) and rabbit polyclonal anti-Ca_V_1.3 (anti-CND1) were kindly provided by Drs. William Catterall and Ruth Westenbroek (University of Washington), and used at 1:250 dilution in blocking solution. Membranes were incubated in primary antibodies overnight at 4°C with constant agitation. The day after, membranes were washed 4x for 10 min in TBST and incubated with Goat anti-rabbit HRP-conjugated secondary antibody at 1:10,000 in blocking solution for 1h at RT with constant agitation. Membranes were washed in TBST, and bands were detected using the Clarity Max Western ECL substrate (Biorad, 1705062) and the iBright imaging system. Protein abundance was calculated by measuring the area under the curve of the band intensity profile using ImageJ (NIH). Each band was normalized to total protein and reported relative to the abundance in young pacemaker samples within the same experiment.

### Electrophysiology

The composition of all the solutions used for electrophysiology can be found in the Solutions section. Recordings of single channels and whole-cell currents were performed at room temperature. Calcium currents were recorded using the whole-cell configuration of the patch-clamp technique in voltage-clamp mode. Isolated pacemaker cells were perfused with Tyrode’s III solution. Borosilicate patch pipettes with resistances of 3–6 MΩ were used. Once the gigaseal was formed, the Tyrode’s III bath solution was exchanged for the I_Ca_ external solution. Current–voltage relationships were obtained by applying a series of 20-ms depolarizing pulses from a holding potential of -75 mV to test potentials ranging from 75 to +75 mV at a 5 mV interval. The voltage dependence of channel activation (G/Gmax) was obtained from the resultant currents by converting them to conductances using the equation, G = I_Ca_/(test pulse potential – reversal potential of I_Ca_); normalized G/Gmax was plotted as a function of test potential. Currents were sampled at 10 kHz and low-pass filtered at 2 kHz using an Axopatch 200B amplifier.

Recordings of action potentials were performed at 32-34°C. Spontaneous pacemaker action potentials were recorded in current-clamp mode using the perforated-patch configuration. On the day of the experiment, freshly prepared β-Escin was added to the intracellular solution to reach a final concentration of 25 μM. Borosilicate patch pipettes with resistances of 3–6 MΩ were used. Spontaneous activity was recorded in the gap-free mode with no holding or transient current applied. Data was sampled at 10 kHz and low-pass filtered at 2 kHz using an Axopatch 200B amplifier. Action potential parameters were analyzed manually using pCLAMP 11 (Molecular Devices).

Single-channel activity was recorded from cell-attached patches. Borosilicate patch pipettes with resistances of 5–7 MΩ were used. Once the gigaseal was obtained, the Tyrode’s III bath solution was exchanged for the single-channel recording external solution. Cells were stimulated with 30 sweeps of a 200 ms square voltage pulse from a resting membrane potential of -80 mV to -30 mV. Current traces were sampled at 10 kHz and low-pass filtered at 2 kHz using an Axopatch 200B amplifier. Currents were analyzed using pCLAMP 11 (Molecular Devices). The single-channel event detection algorithm of pCLAMP was used to measure single-channel opening amplitudes and times. Recordings were filtered using a Gaussian low-pass filter at 1500 KHz, and the baseline was adjusted manually.

### Immunocytochemistry and super-resolution microscopy

For immunostaining Ca_V_1.2 and Ca_V_1.3 channels in pacemaker cells, cells were isolated and plated on PLL-coated coverslips for 2 hours to attach. Excess solution was removed carefully, and cells were fixed with 4% PFA in PBS for 15 min at RT. Unless stated otherwise, all the washing steps in this protocol consisted of 3 rinses with PBS followed by 3 x 5 min washes with PBS on a rocker at low speed. After washing with PFA with PBS, cells were incubated with 50 mM Glycine in PBS for 15 min at RT (aldehyde reduction) and then washed again with PBS. Cells were blocked with 3% w/v BSA - 0.25% v/v Triton X-100 in PBS (blocking solution) for 1h at RT. Primary antibodies were diluted in blocking solution to a concentration of 10 μg/mL (100 μl/coverslip) and incubated overnight at 4°C under gentle orbital agitation. Cells were immunostained using the following antibodies: rabbit polyclonal anti-Ca_V_1.2 (anti-CNC1), rabbit polyclonal Ca_V_1.3 (anti-CND1), guinea pig anti-HCN4 (Alomone, AGP-004). Anti-CNC1 and anti-CND1 were kindly provided by Drs. William Catterall and Ruth Westenbroek (University of Washington). After antibody incubation, cells were washed with PBS and incubated for 1h at RT with specific Alexa Fluor-conjugated antibodies at 2 μg/ml in blocking solution. For Airyscan imaging, Alexa Fluor 647-conjugated donkey anti-rabbit and Alexa Fluor 488-conjugated donkey anti-guinea pig were used. For super-resolution microscopy, Alexa Fluor 647-conjugated donkey anti-rabbit antibodies were used. For super-resolution microscopy, coverslips were mounted on microscope slides with a round cavity (NeoLab Migge Laborbedarf-Vertriebs GmbH, Germany) using MEA-GLOX imaging buffer and sealed with Twinsil (Picodent, Germany). The imaging buffer contained 10 mM MEA, 0.56 mg/ml glucose oxidase, 34 μg/ml catalase, and 10% w/v glucose in TN buffer (50 mM Tris-HCl pH 8, 10 mM NaCl).

A super-resolution ground-state depletion system (SR-GSD, Leica) based on stochastic single-molecule localization was used to generate super-resolution images of Ca_V_1.2 and Ca_V_1.3 channels in pacemaker cells. The Leica SR-GSD system was equipped with 642 nm high-power lasers (2.1 kW/cm2) and a 405 nm 30 mW laser. Images were obtained using a 160X HCX Plan-Apochromat (NA 1.43) oil-immersion lens and an EMCCD camera (iXon3 897; Andor Technology). For all experiments, the camera was running in frame-transfer mode at a frame rate of 100 Hz (10 ms exposure time). Fluorescence was detected through Leica high-power TIRF 642 HP-T filter cube with an emission band-pass filter of 660–760 nm. Super-resolution localization images of Ca_V_1.2 and Ca_V_1.3 channel distribution were reconstructed using the coordinates of centroids obtained by fitting single-molecule fluorescence signals with a 2D Gaussian function using LASAF software (Leica). A total of 50,000 images were used to construct the images.

## Data analysis

Excel (Microsoft), and Prism (GraphPad) were used to analyze data. ImageJ (NIH) was used to process images. Localization maps acquired from super-resolution experiments were reconstructed into .tiff files for processing. Processing consisted of thresholding and binarization of images to isolate labeled structures and analyzed as particles to calculate the area and number. T-test and non-parametric statistics were used to test for significance. p-values <0.05 were deemed statistically significant. The number of cells used for each experiment is detailed in each figure legend. The number of biological samples for each experiment was determined from a power analysis calculation and expertise from previous experiments using the same techniques.

## Acknowledgments

We thank the BRIGHT-UP high-school student, Gabriella Serrano, for her technical support. We thank Dr. William Caterall and Dr. Ruth Westenbroek for providing the anti-L-type calcium channels antibodies used in this study. We thank Dr. Manuel Navedo for his help with the channel coupling analysis. We thank Dr. Bertil Hille and Dr. Rosie Dixon for their critical reading of the manuscript. We thank Dr. Mariya Sweetwyne for the long and fruitful scientific writing sessions.

